# Anti-biofilm efficacy of a medieval treatment for bacterial infection requires the combination of multiple ingredients

**DOI:** 10.1101/2020.04.21.052522

**Authors:** Jessica Furner-Pardoe, Blessing O Anonye, Ricky Cain, John Moat, Catherine A. Ortori, Christina Lee, David A. Barrett, Christophe Corre, Freya Harrison

**Affiliations:** School of Life Sciences, Gibbet Hill Campus, University of Warwick, Coventry, CV4 7AL; Warwick Medical School, Gibbet Hill Campus, University of Warwick, Coventry, CV4 7AL; Warwick Antimicrobial Screening Facility, School of Life Sciences, Gibbet Hill Campus, University of Warwick, Coventry, CV4 7AL; Centre for Analytical Bioscience, Advanced Materials and Healthcare Technologies Division, School of Pharmacy, University of Nottingham, Nottingham NG7 2RD, UK; School of English, University of Nottingham, Nottingham NG7 2RD, UK; Department of Chemistry, University of Warwick, University of Warwick, Coventry, CV4 7AL

## Abstract

Novel antimicrobials are urgently needed to combat the increasing occurrence of drug-resistant bacteria and to overcome the inherent difficulties in treating biofilm-associated infections. Research into natural antimicrobials could provide candidates to fill the antibiotic discovery gap, and the study of plants and other natural materials used in historical infection remedies may enable further discoveries of natural products with useful antimicrobial activity. We previously reconstructed a 1,000-year-old remedy containing onion, garlic, wine, and bile salts, which is known as ‘Bald’s eyesalve’, and showed it to have promising antibacterial activity. In this paper, we have found this remedy has bactericidal activity against a range of Gram-negative and Gram-positive wound pathogens in planktonic culture and, crucially, that this activity is maintained against *Acinetobacter baumannii, Stenotrophomonas maltophilia, Staphylococcus aureus, Staphylococcus epidermidis* and *Streptococcus pyogenes* in a model of soft-tissue wound biofilm. While the presence of garlic in the mixture is sufficient to explain activity against planktonic cultures, garlic alone has no activity against biofilms. We have found the potent anti-biofilm activity of Bald’s eyesalve cannot be attributed to a single ingredient and requires the combination of all ingredients to achieve full activity. Our work highlights the need to explore not only single compounds but also mixtures of natural products for treating biofilm infections. These results also underline the importance of working with biofilm models when exploring natural products for the anti-biofilm pipeline.

**Importance:** Bacteria can live in two ways, as individual planktonic cells or as a multicellular biofilm. Biofilm helps protect bacteria from antibiotics and makes them much harder to treat. Both the biofilm lifestyle and the evolution of antibiotic resistance mean we urgently need new drugs to treat infections. Here, we show that a medieval remedy made from onion, garlic, wine, and bile can kill a range of problematic bacteria grown both planktonically and as biofilms. A single component of the remedy – allicin, derived from garlic – is sufficient to kill planktonic bacteria. However, garlic or allicin alone do nothing against bacteria when they form a biofilm. All four ingredients are needed to fully kill bacterial biofilm communities, hinting that these ingredients work together to kill the bacteria. This suggests that future discovery of antibiotics from natural products could be enhanced by studying combinations of ingredients, rather than single plants or compounds.

## Introduction

Widespread multidrug resistance of once-susceptible pathogens, combined with a lack of success in developing novel antimicrobials, has resulted in a looming crisis. Antimicrobial resistance leads to problematic infections, threatens the success of routine surgery and cancer treatments (1), and is estimated to kill 10 million people per year by 2050 (2,3). One particularly troublesome area is biofilm-associated infection. Biofilm infections are estimated to cost the UK’s National Health Service over a billion pounds every year, with this cost only set to increase (4,5). Biofilms are communities of bacteria that produce a protective extracellular matrix and are especially persistent (6). Biofilm eradication often requires 100-1,000 times higher antibiotic concentrations to achieve clearance than the same bacteria growing planktonically (as individual free-floating cells)(7). *In vivo*, biofilms may essentially be completely impervious to antibiotic treatment (8).

The path to overcoming biofilm infections requires a multifaceted response, including the discovery and clinical deployment of novel antimicrobials. This search for novel candidates must be directed towards those pathogenic bacteria with the highest health and economic impact (e.g. ESKAPE group; (9–11)) and will be expedited if early discovery work on leads takes into account the inherent difficulties of treating biofilm-associated infections (6,12).

The heightened urgency for novel antibiotics has resulted in natural products being revisited. Just 200 years ago, our pharmacopoeia was dominated by herbal medicines. These medicines have had varying levels of success for the treatment of infections in modern research, with a number of potent compounds against planktonic cultures being identified (13,14); however, these have been far less successful against biofilm cultures. We do not fully understand the reasons for this lack of translational success of plant extracts.

It could be that the conventional process for developing drugs may miss key aspects of those herbal remedies which could be effective against biofilms. Conventional drug development calls for the isolation of single active compounds, whereas historical medicine usually calls for combinations of whole plants (and other natural materials). There is some evidence that whole plant extracts can have stronger biological effects than individual isolated compounds (15). Examples of such synergy include observations that the anti-malarial activity of artemisinin is enhanced by the presence of other compounds from the same plant (16,17) or the combination of flavonoids used in Citrox^®^ (15). Synergy may result from the presence of molecules which potentiate the activity of antimicrobial compounds, or from multiple active compounds with different mechanisms of action. It is also possible that in purifying individual compounds to achieve readily quantifiable and characterised treatments, we may lose vital interactions of natural products within the original mixture which prevent irritation or toxicity.

Additionally, research into natural antimicrobial products is often limited by a methodological focus on planktonic activity testing. The current gold standard for antibacterial testing is broth microdilution, usually conducted in cation-adjusted Mueller-Hinton Broth (18), and this is used extensively for the discovery of novel antibacterial compounds. However, whilst providing a great high-throughput screening method, this does not provide information on how the test substance interacts with bacteria in biofilms, or in settings which better mimic the chemical environment experienced by pathogens *in vivo* (18,19). This focus may in part explain the lack of translational success of isolated plant compounds, as their activity may be sufficient to kill planktonic cells, but not to penetrate or kill biofilms – i.e. by not using biofilm assays, we create a high false positive rate in the early stages of activity testing.

In this paper, we investigate the importance of the combination of ingredients for the anti-biofilm activity of a 10th century remedy used for eye infection from the manuscript known as Bald’s Leechbook (London, British Library, Royal 12, D xvii). This remedy has previously been shown to kill *S. aureus* biofilms by Harrison *et al.* (2015) (20), *Pseudomonas aeruginosa* planktonic cultures (21) and recently *Neisseria gonorrhoeae* in a disk diffusion assay (22). The recipe, known as ‘Bald’s eyesalve’, requires equal volumes of garlic (*Allium sativum*) and another *Allium* species (referred to as *cropleac* in the original Old English) to be crushed together and mixed with equal volumes of wine and ox gall (bovine bile). We now report two key new findings. First, we show the bactericidal activity of Bald’s eyesalve against a panel of clinically relevant strains of ESKAPE pathogens in planktonic culture, and against biofilms of a subset of these strains. Second, we conducted a detailed assessment of whether (a) any individual ingredient or (b) the sulphur-containing compound allicin (from garlic) can explain this activity. No single ingredient recaptures the anti-biofilm activity of Bald’s eyesalve, and the presence of allicin cannot fully explain the ability of the whole mixture to kill either planktonic cultures grown in host-mimicking medium (synthetic wound fluid), or biofilms grown in an *in vivo*-like model of a soft tissue wound. The combination of ingredients in the full mixture is the key to the remedy’s apparent efficacy, and this is only apparent when it is tested in host-mimicking models as opposed to planktonic cultures in standard laboratory medium.

## Materials & Methods

### Bacterial strains and culture conditions

Strains used are listed in Table 1. All cultures were grown aerobically at 37°C and Miller’s LB agar (Melford) was used for plating.

**Table 1.**
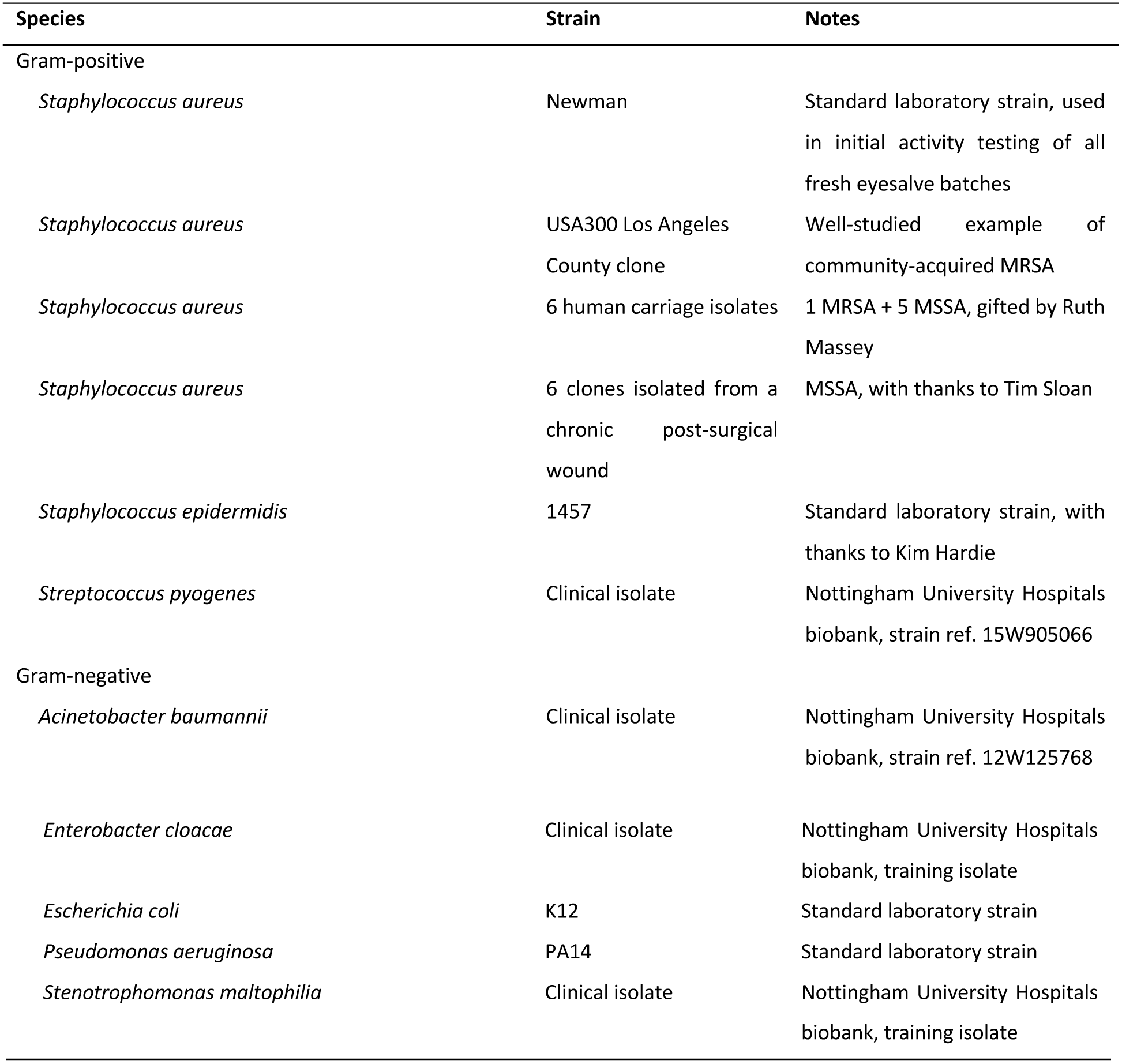
Bacterial strains used in the study

### Media

Synthetic wound fluid (SWF) comprised equal volumes of foetal bovine serum (Gibco) and peptone water (Sigma-Aldrich), (23). Mueller-Hinton Broth (MHB) was obtained from VWR. HyClone water (GE Healthcare) was used throughout.

### Bald’s Eyesalve Reconstruction

Bald’s eyesalve was prepared as previously reported (20). The outer skin of the garlic and onion (sourced from local greengrocers) was removed. The garlic and onion were finely chopped, and equal volumes of garlic and onion were crushed together using a mortar and pestle for 2 minutes. Various sized batches of Bald’s eyesalve were used throughout this paper, ranging from final volumes of 30 ml – 400 ml, the average weight used was 14.1 ± 1.5 g of onion and 15.0 ± 1.3 g of garlic per 100 ml of Bald’s eyesalve.

The crushed onion and garlic were then combined with equal volumes of wine (Pennard’s organic dry white, 11% ABV, sourced from Avalon Vineyard, Shepton Mallet) and bovine bile salts (Sigma Aldrich) made up to 89 mg·ml^-1^ in water and sterilised by exposing to UV radiation for 10 minutes (Carlton Germicidal Cabinet fitted with a 2537Å, 8-Watt UV tube). The mixture was stored in sterilised glass bottles in the dark at 4°C for 9 days, after which it was strained and centrifuged for 5 minutes at 1,811 g. The supernatant was then filtered using Whatman™ 1001-110 Grade 1 Qualitative Filter Paper, Diameter: 11 cm, Pore Size: 11 µm. Filtered Bald’s eyesalve was stored in sterilised glass vials in the dark at 4°C.

For ease of reading, batches are numbered as they appear within the paper. The key for their batch name is provided in the Data Supplement.

### Preparation of Individual Ingredients and Dropouts

Individual ingredients were prepared such that their concentrations were equal to concentrations present in the full remedy, a schematic is provided in Figure S1. To achieve comparable concentrations, each ingredient was prepared as it would be for the full remedy (onion and garlic crushed separately from each other), with all other ingredients substituted with the same volume of water. A similar process was followed for “dropout” batches, where one ingredient at a time was systematically excluded from the remedy and replaced with an equal volume of water to maintain comparable concentrations to the original remedy.

### Planktonic killing assay

Bacteria were cultured aerobically at 37°C on LB agar for 18-24 hours. Several colonies were then inoculated into 5 ml SWF and incubated for 6 hours at 37°C on an orbital shaker. Aliquots of each bacterial culture (100 µl) were added to wells of Corning^®^ Costar^®^ TC-Treated 96-Well Plate and 50 μL of the eyesalve batch to be tested was added to 5 wells per strain. 50 µL of water was used as a negative control. Plates were incubated in Tecan SPARK 10M at 37°C for 18 hours with periodic orbital shaking (10 seconds at 20 min intervals). Serial dilutions were then performed and plated on LB agar plates.

### Minimum Inhibitory Concentration Assay by Broth Microdilution

The minimum inhibitory concentrations (MIC) of eyesalve, single ingredients, dropout batches and allicin were determined as described by Wiegand, Hilpert and Hancock (2008) in MHB and in SWF. Treatments, or water as a negative control, were serially diluted with media in Corning^®^ Costar^®^ TC-Treated 96-Well Plates, following the scheme in Table 2. Bacterial isolates were streaked onto LB agar to obtain single colonies. After 18 - 24 hours incubation at 37°C, three to five morphologically similar colonies were transferred to phosphate-buffered saline (PBS) and diluted to 0.5 McFarland standard (OD_600_ 0.08-0.1). This suspension was diluted 1 in 100 in media, resulting in 5 × 10^5^ CFU/ml. The bacterial suspension was then added to the treatment dilutions, resulting in another twofold dilution. The highest final concentration of eyesalve, single ingredient or dropout batch tested was 50%. Plates were incubated for 18 hours at 37°C. Results were visually checked for turbidity, and MIC values are the lowest concentration where growth was not visible.

**Table 2.** Final volumes and calculated percentages of Bald’s eyesalve in wells of 96-well plates used for MIC.

| Well | Bald's eyesalve (μl) | Media (μl) | Bacterial suspension (μl) | Final volume (μl) | % ESO in well |
| --- | --- | --- | --- | --- | --- |
| **A** | 50 | 0 | 50 | 100 | 50 |
| **B** | 33 | 17 | 50 | 100 | 33 |
| **C** | 25 | 25 | 50 | 100 | 25 |
| **D** | 12.5 | 37.5 | 50 | 100 | 12.5 |
| **E** | 6.25 | 43.75 | 50 | 100 | 6.25 |
| **F** | 3.13 | 46.87 | 50 | 100 | 3.13 |
| **G** | 1.56 | 48.44 | 50 | 100 | 1.56 |
| **H** | 0.78 | 49.22 | 50 | 100 | 0.78 |

### Biofilm Killing Assay in Synthetic Wounds

Biofilms were created in a synthetic soft-tissue wound model as described by Werthén *et al.* (2010; (23)) and as used in our previous work (20). Briefly, synthetic wounds were created on ice and comprised 2 mg·ml^-1^ collagen, 0.01% acetic acid, 60% [vol/vol] SWF, 10 mM sodium hydroxide. Synthetic wounds were incubated at 37°C for 1 hour to allow collagen to polymerise and placed under UV light for 10 minutes to ensure no contamination. Wounds were either 400 µl in 24-well culture plates or 200 µl in 48-well plates.

Bacteria were incubated aerobically in SWF on an orbital shaker for 6 hours at 37°C, after which cultures were diluted to OD_600_ of 0.1 - 0.2 with fresh SWF. Bacteria suspension was added to each synthetic wound (100 µl in 400 µl wounds or 50 µl in 200 µl wounds) and incubated at 37°C for 24 hours to allow biofilm formation.

Wounds containing mature biofilms were then exposed to Bald’s eyesalve or water (200 µl in 400 µl wounds or 100 µl in 200 µl wounds). Bacteria were recovered by treating with 300 μl (400 µl wounds) or 150 µl (200 µl wounds) of 0.5 mg·ml^-1^ collagenase type 1 (EMD Millipore Corp, USA) for 1 hour at 37°C to break down the matrix; the resulting liquid was serially diluted and plated on LB plates. Plates were incubated at 37°C overnight, colonies were counted, and CFU/wound calculated.

### High-Performance Liquid Chromatography Quantification of Allicin

Allicin is an unstable and reactive compound under gas chromatography settings (24), therefore, high-performance liquid chromatography (HPLC) was used to determine allicin concentrations (24).

Various preparations of Bald’s eyesalve were analysed by reversed-phase HPLC on an Agilent 1200 series system fitted with an Agilent ZORBAX Eclipse XDB-C18, 150 x 4.6 mm, 5 µm particle size column and diode array detection at 210 nm. A gradient of methanol (5-95%) in water was used at a flow rate of 1 ml·min^-1^ over 30 minutes. Injection volume was 10 μl, and the column temperature was 25°C.

To identify the concentration of allicin in eyesalve preparations, a calibration curve was prepared by serially diluting an external allicin standard (Abcam, Cambridge) in water (twofold dilutions, from 750 µg·ml^-1^ to 0.5 µg·ml^-1^). External standards were run through HPLC as described above (n = 2) and allicin peak area, in the 210 nm reading, was plotted against concentration (µg·ml^-1^) (Figure S2A and Data Supplement). Allicin external standards had a retention time of approximately 15 minutes (Figure S2B). This peak was confirmed in Bald’s eyesalve by comparing fresh eyesalve to the same batch spiked with additional allicin standard. The suspected peak increased as expected (Figure S2C-D).

### Statistics

Data were analysed in R v3.5.1 (R Core Team, 2018) using the *car* (25), *lsmeans* (26), *multcomp* (27) and *FSA* (28) packages. Raw data and R code for all experiments are provided in the Data Supplement.

## Results

### Consistent anti-biofilm activity of Bald’s eyesalve and decision to use onion (Allium cepa) as “cropleac”

The meaning of Old English *cropleac*, as used in the original remedy, is ambiguous and may refer to a variety of *Allium* species (29,30). Two likely translations are onion (*Allium cepa*) or leek (*Allium porrum*). For this reason, multiple batches of the remedy using either *Allium* species were made in our laboratory between 2014 and 2019. After the nine-day brewing period, the activity of the remedies was tested against mature biofilms of *S. aureus* Newman in an *in vivo*-like model of a soft tissue wound. In this assay, 24-hour-old biofilms, that have been grown at 37°C in collagen-based synthetic wounds, are exposed to Bald’s eyesalve or water for 24 hours, and surviving bacteria are counted (20,23).

In total, 75 batches of Bald’s eyesalve were made, including 15 pairs of batches where onion (ESO) and leek (ESL) variants were made at the same time, using the same garlic, wine and bile. As shown in Figure 1A, biofilm killing was achieved by 14/15 paired batches, with 22/30 preparations causing a >3-log drop in viable bacteria, when compared with water-treated control biofilms. The mean log drop was not significantly different between paired ESO and ESL batches (paired *t*-test t_14_= 0.025, *p* = 0.981). We tested one example pair of ESO and ESL variants (batch 6) against a panel of twelve clinical isolates of *S. aureus* grown in the synthetic wound biofilm model. As shown in Figure 1B, ESO consistently showed more bactericidal activity than ESL. This result may be due to the presence of antimicrobial compounds unique to onion, or it may simply be due to onions being easier to crush in a mortar and pestle, potentially making the extraction of natural compounds more efficient. To simplify further analysis, follow on work was focussed on ESO.

**Figure 1.**
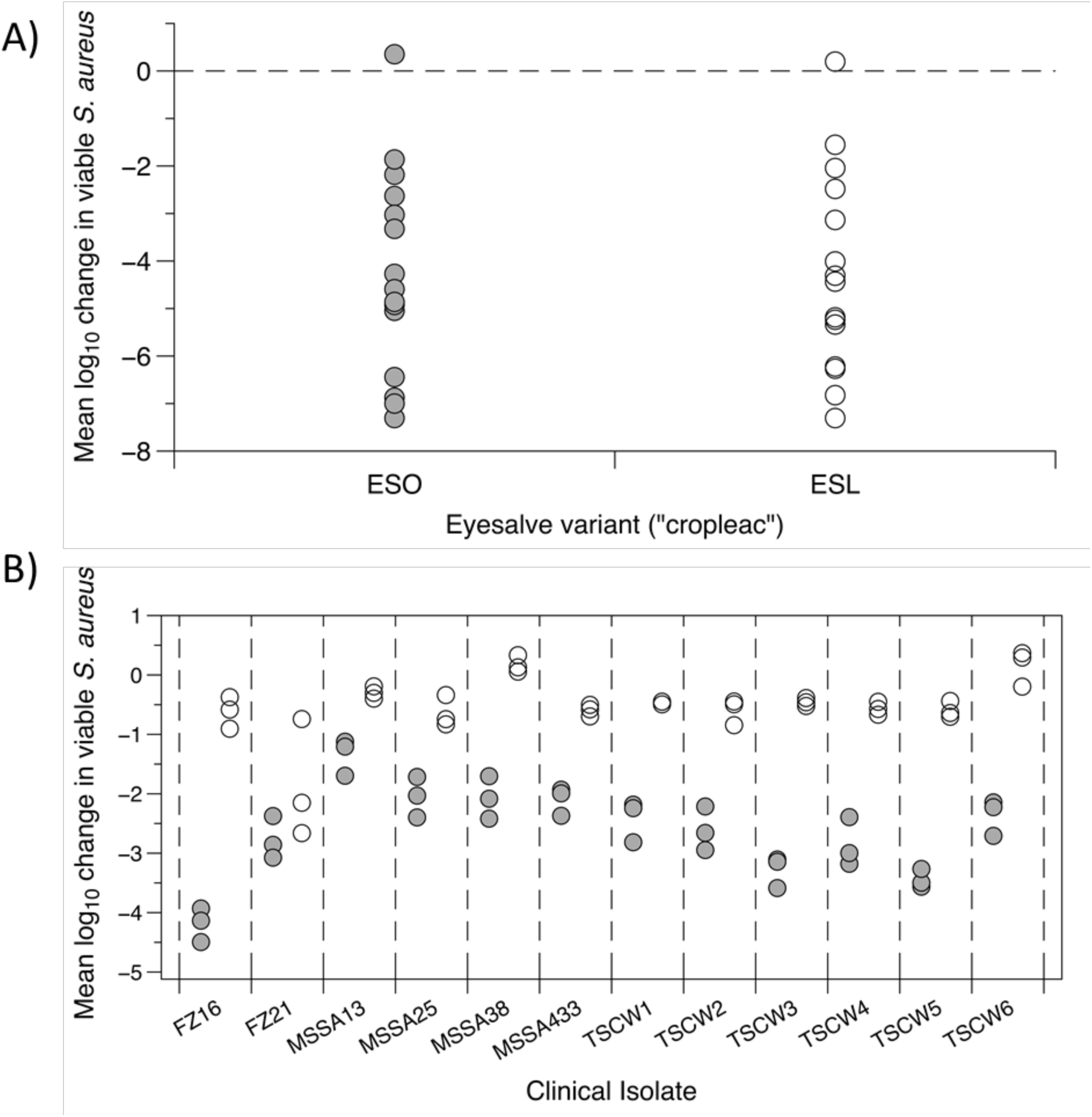
The activity of Bald’s eyesalve, ESO and ESL variants, against *S. aureus*. (A) 15 pairs of Bald’s eyesalve translating *cropleac* as either onion (ESO, grey) and leek (ESL, white), were prepared at the same time (batches 1-15). Their activity was assessed against *S. aureus* Newman biofilms. Mature biofilms of *S. aureus* Newman were grown in a model of a soft tissue wound, then treated with either sterile water or Bald’s eyesalve (n = 3-5 replicates per treatment) for 24 hours before recovering bacteria for CFU counts. The mean log change in viable bacteria in treated vs control wounds was calculated for each preparation and no significant difference was seen (paired *t*-test t_14_= 0.025, *p* = 0.981). Raw data are supplied in the Data Supplement. (B) Six carriage isolates of *S. aureus*, and six isolates from a chronic post-surgical wound (detailed in Table 1), were grown in the synthetic wound biofilm model and treated with ESO, ESL or water (n=3 per treatment). Each strain grew to different densities in untreated wounds (range approx. 10^6^–10^8^). The log_10_ drop in bacteria associated with treatment was calculated for each treated wound relative to the mean CFU in the three untreated wounds. Log drop data was analysed by ANOVA which revealed significant differences in log drop between eyesalve variants (F_1,48_ = 732, *p* < 0.001) and strains (F_11,48_ = 14.4, *p* < 0.001) and a significant strain-dependent difference in the magnitude of the effect of eyesalve variant (strain*variant interaction F_11,48_ = 8.66, *p* < 0.001). The family-wise error rate was calculated from the ANOVA table and used to conduct planned contrasts of ESO vs ESL treatment effects for each strain using *t-*tests. ESO caused a larger reduction in viable cells than ESL for all strains (all *p* < 0.001). Raw data and R scripts are supplied in the Data Supplement.

As shown in Figure 2, biofilm killing was consistently achieved by multiple batches of ESO. Of these batches, 62 of the 75 caused a >3-log drop in viable CFU, with 45/75 causing >5 log drop, and 28 of these causing > 6-log drop, all relative to control biofilms treated with sterile water.

**Figure 2.**
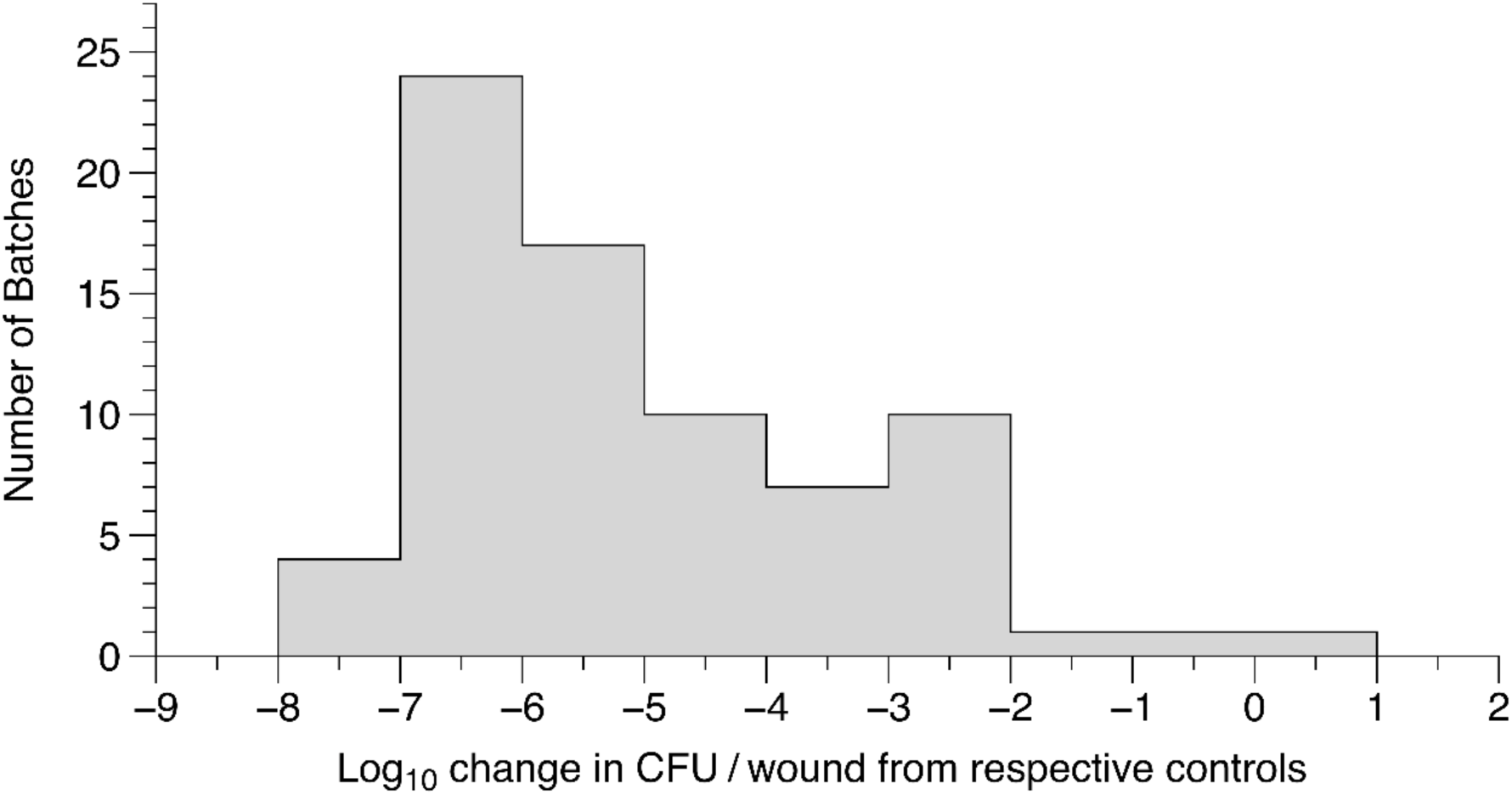
The activity of 75 Bald’s eyesalve (ESO) batches made by our group against *S. aureus* Newman biofilms. Mature *S. aureus* Newman biofilms were grown in a model of a soft tissue wound, then treated with either sterile water (control) or eyesalve (n = 3-5 replicates per treatment) for 24h before recovering bacteria for CFU counts. All batches showed killing, and the log drop in treated vs control wounds was calculated. Raw data are provided in the Data Supplement.

### Broad-spectrum antibacterial activity of Bald’s eyesalve against common wound pathogens

The antibacterial activity of three batches of ESO was tested in both planktonic and biofilm cultures of eight strains of bacteria that commonly cause chronic wound infections. As shown in Figure 3, ESO had potent activity against planktonic cultures of Gram-negative (*P. aeruginosa* PA14, *A. baumannii* clinical isolate, *E. cloacae, S. maltophilia*) and Gram-positive (*S. aureus* Newman, *S. aureus* USA300, *S. epidermidis* and *S. pyogenes*) wound pathogens. ESO eradicated all planktonic cultures in all strains tested with the exception of *S. aureus* USA300 and *S. maltophilia*, where a 3-4 log drop in viable bacteria was seen.

**Figure 3.**
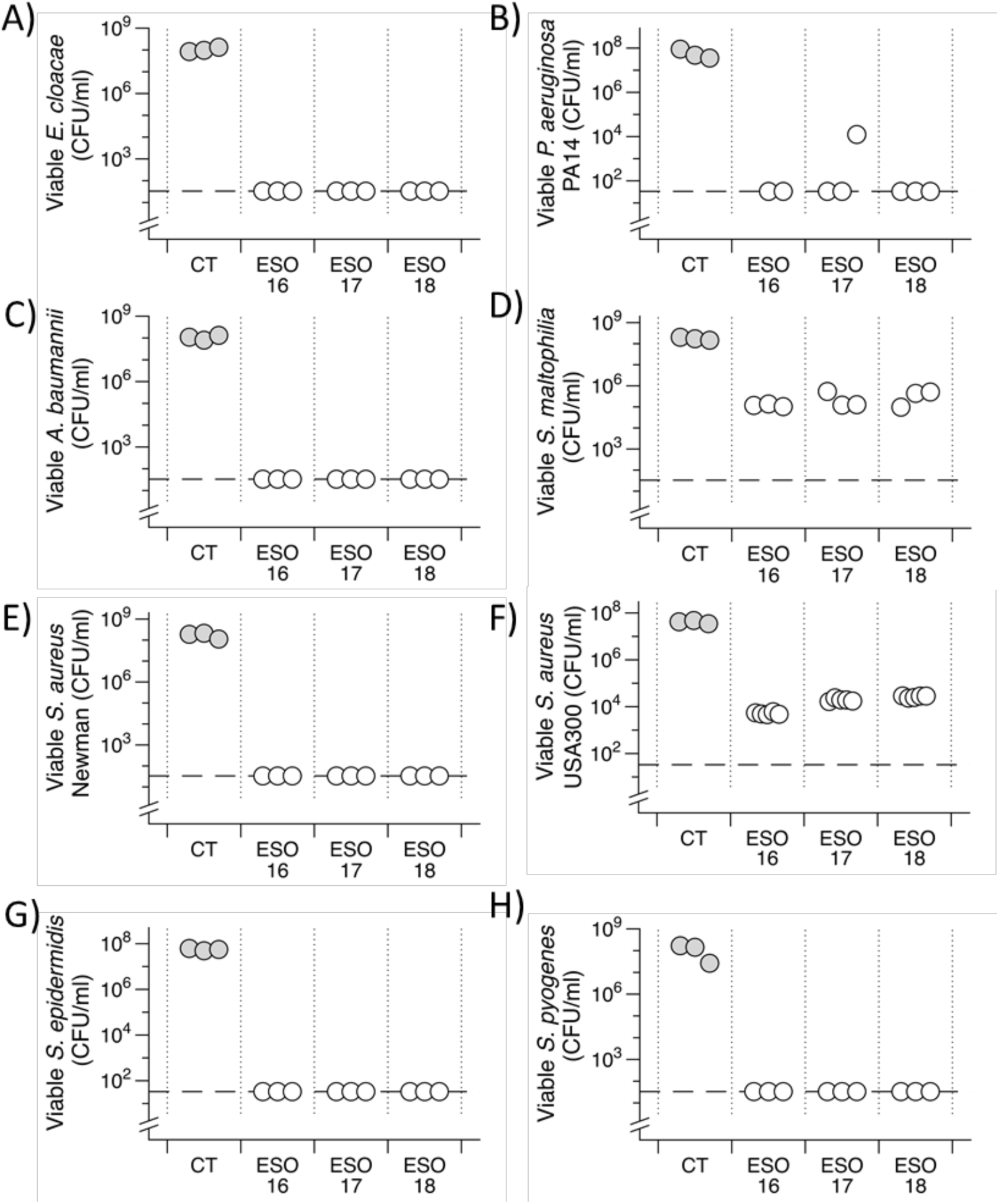
Bald’s eyesalve (ESO) activity against eight example strains of bacteria that commonly cause chronic wound infections. Planktonic cultures were grown for 6 hours in synthetic wound fluid. Cultures were treated with ESO (batches 16-18) or sterile water (CT) to a final concentration of 33% v/v (n = 3-5 replicates per treatment). After 18 hours bacteria were recovered for CFU counts. The dashed line represents the limit of detection by plating. For *S. maltophila* and *S. aureus* USA300, where we did not observe complete killing, we used ANOVA to determine that the CFU recovered from ESO-treated wounds was significantly different from control wounds (*S. aureus* USA300: F_2,14_ = 3458, *p* < 0.001; *S. maltophilia*: F_3,8_ = 87.41, *p* < 0.001). Data were log transformed to meet the assumptions of linear modelling. A Dunnett’s test was conducted to compare the CFU of the three batches with the CFU of the controls. All 3 batches of ESO were significantly different from their respective controls for both isolates (all *p* < 0.001). Raw data and R scripts are supplied in the Data Supplement.

Mature biofilms of the above isolates were grown in the synthetic wound model and treated with the same batches of eyesalve used for planktonic killing experiments. A 2-6-log drop in viable cells was observed for the Gram-positives *S. aureus* Newman, *S. aureus* USA300, *S. epidermidis, S. pyogenes* and the Gram-negative *A. baumannii* (Figure 4). No or inconsistent killing was observed for *P. aeruginosa, E. cloacae* and *S. maltophilia* biofilms (Figure S3).

**Figure 4.**
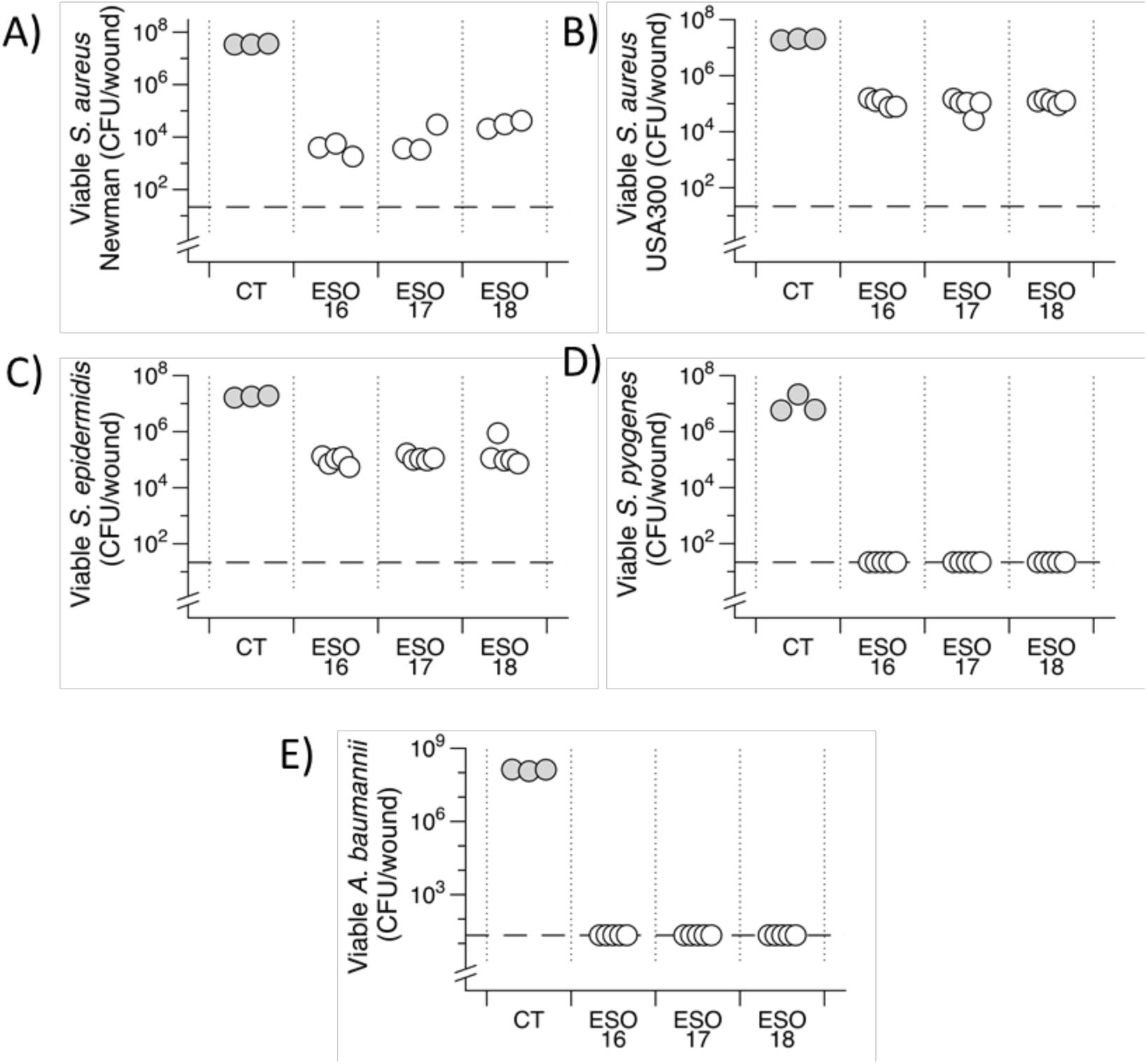
Bald’s eyesalve (ESO) anti-biofilm activity. Mature biofilms of various isolates were grown in a model of a soft tissue wound, then treated with either sterile water (control, CT) or eyesalve (batches 16-18), to a final concentration of 33% (v/v), for 24h before recovering bacteria for CFU counts (n = 3-5 replicates per treatment). The dashed line represents the limit of detection by plating. For *S. aureus* Newman, *S. aureus* USA300, and *S. epidermidis*, where we did not observe complete killing, we used ANOVA to determine that the CFU recovered from ESO-treated wounds was significantly different from control wounds (*S. aureus* Newman: F_3,8_ = 107, *p* < 0.001; *S. aureus* USA300: F_3,14_ = 103.5, *p* < 0.001; *S. epidermidis*: F_3,14_ = 63.86, *p* < 0.001). Data were log transformed to meet the assumptions of linear modelling. A Dunnett’s test was conducted to compare the CFU of the three batches with the CFU of the controls. All 3 batches of ESO were significantly different from their respective controls for all three isolates (all *p* < 0.001). Raw data and R scripts are supplied in the Data Supplement.

### Garlic is responsible for the majority of planktonic killing by ESO

Our initial work indicated that all four ingredients in ESO were required to kill biofilms of *S. aureus* Newman in synthetic wound biofilms (20). A recent publication by Fuchs *et al.* (2018; (21)) concluded that the bactericidal activity of ESO was due to the presence of allicin from garlic, however, the Fuchs *et al.* study only investigated planktonic killing in standard Mueller-Hinton Broth (MHB). It is well known that planktonic cultures of bacteria can be up to 1,000 times more sensitive to antibiotics than the same isolates grown as biofilms (7), and that antibiotic sensitivity is highly dependent on growth medium (31). This means conducting assays only on planktonic MHB cultures risks underestimating the number or concentration of bioactive agents in ESO required for killing in *in vivo*-like conditions (18,19).

To more robustly determine if the activity of ESO stems from one ingredient or several, we prepared individual ingredients and preparations omitting one ingredient, such that the concentrations of each ingredient in the whole remedy and the single ingredient or dropout variants were equal. The MICs of these preparations, along with MICs of the full recipe, were assessed in MHB and in SWF, in standard broth microdilution assays using four of the isolates previously tested (two Gram-negatives, *A. baumannii* clinical isolate strain and *P. aeruginosa* PA14, and two Gram-positives, *S. aureus* Newman and *S. aureus* USA300).

As shown in Table 3, the bacterial isolates varied in their sensitivity to the full ESO. Crucially, there was an effect of the growth medium, such that MICs at least doubled in SWF compared with MHB for *S. aureus* Newman and *A. baumannii*. Wine, bile or onion alone were much less effective than the full remedy. In MHB, the MIC of garlic alone was the same as the ESO MIC for all strains except *P. aeruginosa*, and in SWF the garlic and ESO MICs were either equal or exhibited a max. 2-fold difference. This suggests that the planktonic activity of the remedy is due to the presence of garlic. The results of MIC testing with ingredients omitted gave similar results (Table 4). Preparations omitting garlic lost all or most of their antibacterial activity. Removal of any other ingredient left activity largely unaffected, although the omission of wine or bile doubled the MIC against *P. aeruginosa* PA14 in SWF. The discordance between MICs in different media is more prominent here, with the removal of an ingredient having a minimal effect in MHB but a large loss in activity when tested in SWF. This stresses the common discordance between testing in standard rich lab medium and host-mimicking medium.

**Table 3.**
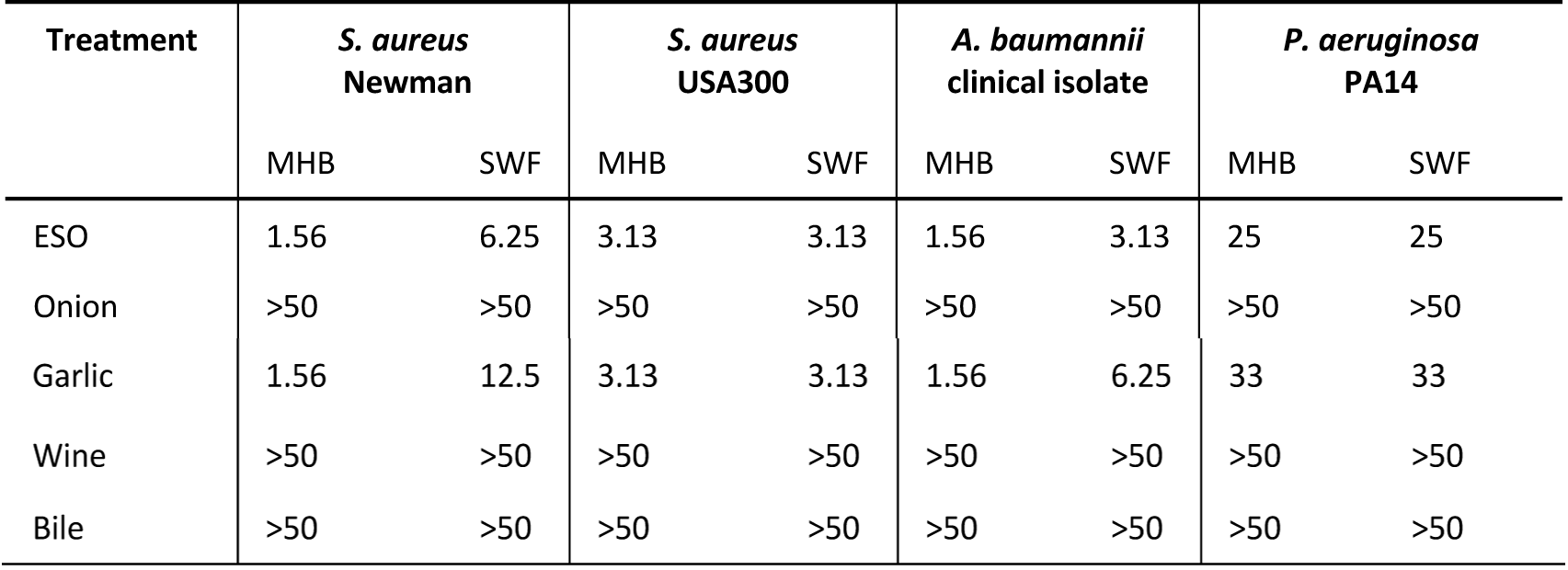
Minimum inhibitory concentration (MIC) of ESO or individual ingredients. MICs were tested in both Mueller-Hinton Broth (MHB) or synthetic wound fluid (SWF). MICs are presented as modal values of 3 different batches (ESO 19-21; with 3 replicate MIC tests per batch) and are the percentage of treatment present at MIC (v/v).

|  |  |  |  |  |  |  |  |  |
| --- | --- | --- | --- | --- | --- | --- | --- | --- |
| Garlic | 1.56 | 12.5 | 3.13 | 3.13 | 1.56 | 6.25 | 33 | 33 |
| Wine | >50 | >50 | >50 | >50 | >50 | >50 | >50 | >50 |
| Bile | >50 | >50 | >50 | >50 | >50 | >50 | >50 | >50 |

**Table 4.**
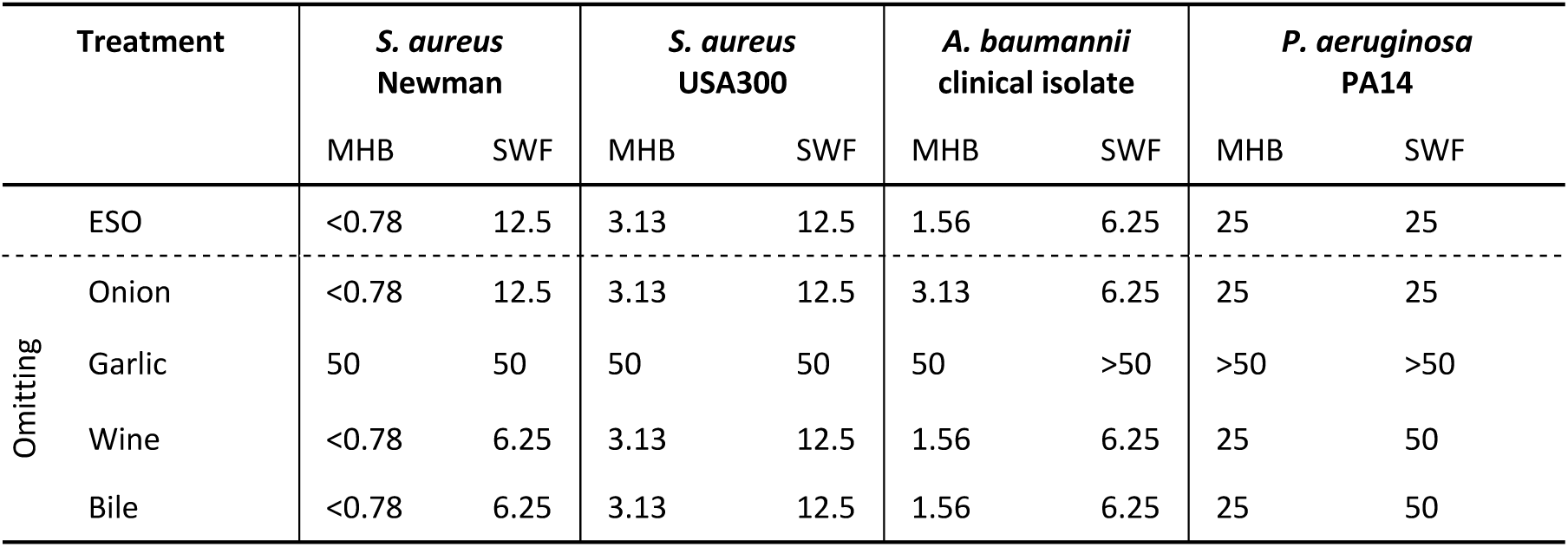
Minimum inhibitory concentration (MIC) of ESO or preparations omitting a single ingredient. MICs were tested in both Mueller-Hinton Broth (MHB) and synthetic wound fluid (SWF). MICs are presented as modal values of 3 different batches (ESO 22-24; with 3 replicate MIC tests per batch) and are the percentage of treatment present at MIC (v/v).

### All four ingredients are necessary for activity against mature *S. aureus* biofilms in synthetic wounds

Consistent with previous results from our group, no single ingredient alone had any bactericidal activity against mature *S. aureus* Newman biofilms in the synthetic wound model, and removing any single ingredient resulted in a reduction in the anti-biofilm activity (Figure 5). Surprisingly, removal of wine caused a decrease in activity on par with that seen with the removal of garlic, despite wine possessing very limited antimicrobial activity on its own in either biofilm or planktonic assays (Table 3). For comparison with planktonic MIC data, the final concentration of Bald’s eyesalve in synthetic wound biofilm killing assays (Figures 1-6), is 33% vol/vol. Therefore, in more *in vivo*-like biofilm conditions, combining all the ingredients is necessary for full activity against *S. aureus* Newman.

**Figure 5.**
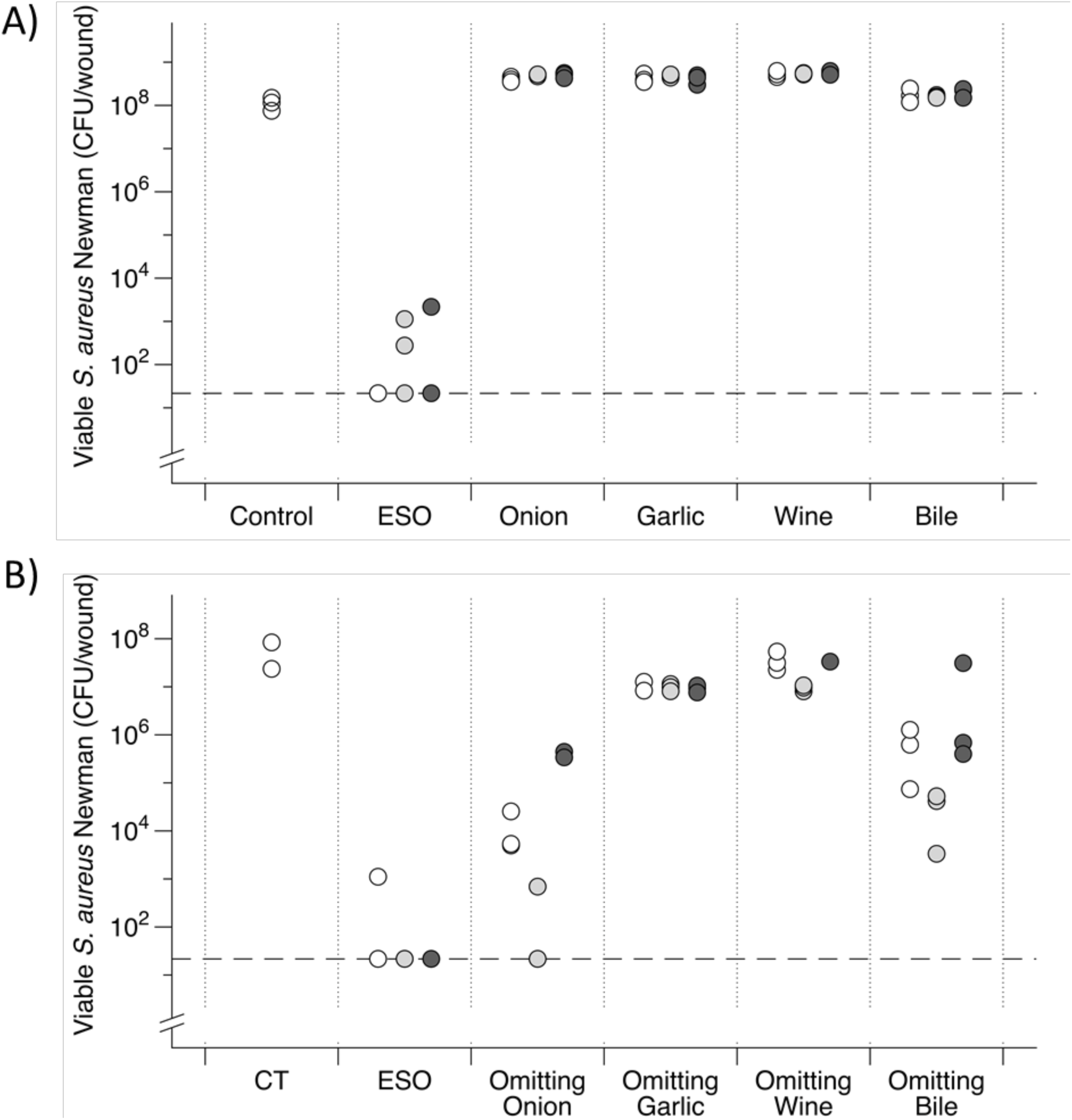
Contribution of each ingredient to Bald’s eyesalve (ESO) anti-biofilm activity. Mature biofilms of *S. aureus* Newman were grown in a model of a soft tissue wound, then treated with either water (CT) or treatment for 24 hours, before recovering bacteria for CFU counts. The dashed lines represent the limit of detection by plating. A) ESO and individual ingredients prepared such that the concentrations of each ingredient in the whole remedy and individual preparations are equal (n = 3 replicates per treatment). Three independent batches of preparations are shown, corresponding to batches ESO 19, 20 and 21, shown in white, light grey and dark grey, respectively. Analysis of the data by ANOVA showed a significant effects of treatment (F_4,30_ = 389.215, *p* < 0.001) but no effect of batch (F_2,30_ = 1.24, *p* = 0.305) and no interaction between treatment and batch (i.e. the effect of treatment did not depend on batch: F_8,30_ = 0.930, *p* = 0.507). Data were square root transformed to meet the assumptions of linear modelling. To compare each treatment group to the untreated controls, we then fitted a new ANOVA to fit least-squares means and variances for viable CFU recovered after each treatment (excluding controls), averaged over the three batches. These fitted means were then compared to the mean observed in the untreated controls using a two-tailed, unpaired Welch’s *t*-test, which takes into account the unequal sample sizes and variances in control and treatment groups. The decrease in CFU on treatment with full ESO was significant (*t*_2.93_ = 9.18, *p* = 0.003). Garlic, onion and wine caused small but significant increases in viable CFU (*t*_2.93_ = 9.05, *p* = 0.003, *t*_2.93_ = 9.61, *p* = 0.003, *t*_2.93_ = 11.1, *p* = 0.002), while bile had no significant effect on CFU (*t*_2.93_ = 2.44, *p* = 0.094). B) ESO and batches with one ingredient omitted and replaced with water, to keep the concentrations of remaining ingredients equal to those in the full recipe. Three independent batches of preparations are shown, corresponding to ESO batches 22, 23 and 24, shown in white, light grey and dark grey, respectively. ANOVA showed significant differences in CFU between treatments (F_4,26_ = 162, *p* <0.001) and between batches (F_2,26_ = 18.7, *p* <0.001), and a significant interaction between treatment and batch (F_8,26_ = 5.85, *p* <0.001); the graph shows that this interaction is due to different batches showing larger or smaller differences between the killing effects of the preparations, but that the rank order of preparations by CFU is the same in each batch. Data were log-transformed to meet the assumptions of linear modelling. As only two control data points were obtained for this experiment, one-sample *t-*tests were performed using least-squares mean CFU in the treated wounds to the mean CFU in control wounds; the decrease in CFU vs control was statistically significant for ESO and all dropouts (*p* ≤ 0.002) except the wine dropout (*p* = 0.202). A Dunnett’s test was conducted to compare each dropout with the full ESO. All dropouts resulted in higher CFU than the full ESO (*p* < 0.001). Raw data and full statistical results are supplied in the Data Supplement.

**Figure 6.**
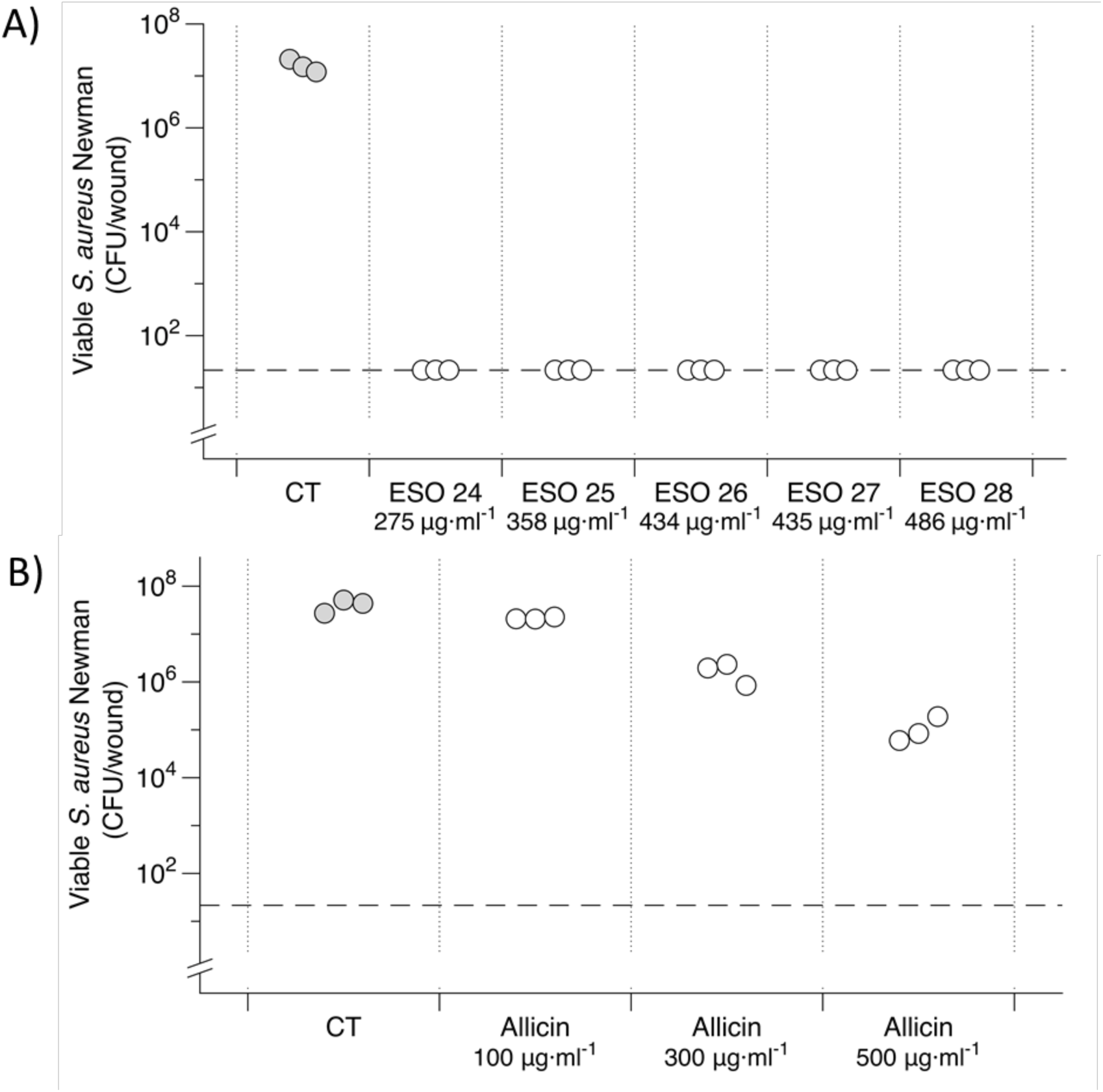
Antimicrobial activity of Bald’s eyesalve (ESO) and allicin standards. Mature *S. aureus* Newman biofilms were grown in a model of a soft tissue wound, then treated with either water (CT) or treatment for 24 hours, before recovering bacteria for CFU counts. A) Biofilms treated with 5 ESO batches, with their respective allicin concentrations indicated. B) External allicin standards diluted in water. The dashed line represents the limit of detection by plating. To statistically analyse, data were log transformed to meet the assumptions of linear modelling. A one-way ANOVA was performed and found a significant difference between allicin treatments and control cells (F_3,9_ = 141, *p* < 0.001). A Dunnett’s test was conducted to compare the CFU of the three allicin concentrations with the CFU of the controls. Of the concentrations, 300 µg·ml^-1^ and 500 µg·ml^-1^ were significantly different (*p* < 0.001), 100 µg·ml^-1^ was not significantly different (*p* = 0.247). Raw data and full statistical results are supplied in the Data Supplement.

### The antibacterial activity of ESO is not simply due to its allicin content

The study which attributed the planktonic activity of ESO to garlic further concluded that this was specifically due to the presence of allicin (21), an organosulfur compound with well known, potent antimicrobial activity *in vitro* (32–34). We used HPLC, calibrated against allicin standards of known concentrations, to determine the allicin concentration in 12 batches of fresh ESO. These batches were then assayed for MIC against *S. aureus* Newman in SWF. We were then able to calculate the concentration of allicin present in the MIC of each batch and compare this with the MIC of purified allicin.

As shown in Table 5, the allicin concentration in the 12 batches of ESO was found to be in the range of 391 ± 19 µg·ml^-1^. Therefore, the estimated concentration of allicin in their respective MICs would be in the range 11 - 39 µg·ml^-1^. This is much lower than the MIC obtained for purified allicin, which was 62.5 µg·ml^-1^. Due to experimental error, two MIC test (ESO batches 28, 29) were lost; we did not repeat this due to the time delay between the initial and repeated test potentially affecting concentrations of compounds present, and thus activity.

**Table 5.** Minimum inhibitory concentrations (MIC) were calculated for fresh Bald’s eyesalve (n = 12; batches 19-30), garlic only preparations (n = 3) and external allicin standards (n = 3). Allicin concentrations for each batch were quantified using HPLC and calculated using an allicin calibration curve. MICs were measured in synthetic wound fluid. The allicin concentration in the MIC of each batch, and the allicin concentration present in the biofilm treatment experiments, is calculated and the results of biofilm treatment experiments are provided for reference. Raw data is supplied in the Data Supplement.

| Treatment | Allicin concentration range<br>(mean ± s.e.) | Allicin<br>concentration<br>in MIC | Biofilm activity (log<br>drop from control) | Average predicted<br>allicin concentration<br>in biofilm treatment |
| --- | --- | --- | --- | --- |
| ESO | 275 - 486 ($391 \pm 19$) $\mu\text{g}\cdot\text{ml}^{-1}$ | 11 - 39 $\mu\text{g}\cdot\text{ml}^{-1}$ | Eradicated biofilm (>5) | 142 $\mu\text{g}\cdot\text{ml}^{-1}$ |
| Garlic only | 358 - 434 ($389 \pm 19$) $\mu\text{g}\cdot\text{ml}^{-1}$ | 45 - 54 $\mu\text{g}\cdot\text{ml}^{-1}$ | No activity (0) | 160 $\mu\text{g}\cdot\text{ml}^{-1}$ |
| Allicin<br>Standard | | 62.5 $\mu\text{g}\cdot\text{ml}^{-1}$ | Partial activity (2.6) | 167 $\mu\text{g}\cdot\text{ml}^{-1}$ |

All of these 12 batches eradicated mature *S. aureus* Newman biofilms in the synthetic wound assay (see Data Supplement). The results of biofilm killing assays for five representative batches are depicted in Figure 6A, alongside a biofilm killing assay conducted for preparations of 100, 300 or 500 µg·ml^-1^ of purified allicin. Purified allicin at concentrations similar to those found in ESO, had much less anti-biofilm activity (Figure 6B), and garlic preparations with the same allicin concentration as ESO had no activity (data supplied in the supplementary data file). Therefore, allicin alone does not explain either the activity of ESO against biofilm-grown bacteria or its activity against planktonic cultures.

## Discussion

The activity of a range of natural products against microbes in simple *in vitro* assays (agar diffusion, broth microdilution or simple surface-attached biofilm assays) demonstrates their potential as a source of novel antibiotics (35–37). However, the standard approach of purifying individual compounds from natural products rarely produces clinically useful products and potent activity against planktonic bacteria in standard lab media rarely translates to *in vivo* efficacy. This is an especially pressing problem in the case of biofilm infections. Biofilms are much harder to treat due to reduced penetration of antibiotics through the extracellular matrix and the enhanced tolerance of biofilm-grown cells to many in-use antibiotics (6,12). Biofilm infections of wounds (e.g. burns, diabetic foot ulcers), medical implants (e.g. artificial joints, catheters), the lungs (e.g. in cystic fibrosis) and other body sites impose a major health and economic burden (38) and can be effectively untreatable. Non-healing, infected foot ulcers, which can be a complication of diabetes, provide an especially sobering example. Even if the infection is apparently successfully treated, there is a high chance of recurrence and an estimated 50% of those affected die within five years of ulcer development. Management of diabetic foot ulcers costs the UK’s NHS £650M per year (39).

Historical medical manuscripts often prescribe complex preparations of several ingredients to treat infections. Thus, when considering natural products as a potential source of anti-biofilm agents, we must consider the possibility that any efficacy they may possess could rely on creating a cocktail of different products. Understanding the relationship between combinations of natural products and antimicrobial activity may generate a novel way to create new antibiotics from botanicals. Here, we confirm Bald’s eyesalve as an example of an “ancientbiotic” that requires the combination of all ingredients for potent activity against a panel of clinically important bacterial strains.

Our research builds on previous work (20–22) to show that Bald’s eyesalve can eradicate planktonic cultures of a range of problematic Gram-positive and Gram-negative bacteria including *P. aeruginosa, A. baumannii, E. cloacae, S. maltophilia, S. aureus, S. epidermidis* and *S. pyogenes*. It is also able to cause a 4-log reduction in viable cell counts in planktonic cultures of the especially problematic MRSA. Despite the widely understood problems of treating biofilms (6,12), Bald’s eyesalve was also able to significantly reduce viable cell counts in biofilms of *S. epidermidis* and MRSA and was able to completely eradicate biofilms of *S. aureus* Newman, *A. baumannii* and *S. pyogenes*, in an established soft-tissue wound model (23). However, although there was promising planktonic activity against *P. aeruginosa, E. cloacae* and *S. maltophilia*, variable or no activity was seen against biofilm cultures of these isolates in the wound model. This highlights the importance of investigating the anti-biofilm activity of candidate antibacterial agents, rather than extrapolating from the results of planktonic assays. Bald’s eyesalve shows great promise as an effective antimicrobial candidate, although further development and combination with biofilm-degrading adjuvants may be necessary to achieve activity against species such as *P. aeruginosa*.

Each of Bald’s eyesalve ingredients has known antimicrobial properties or compounds (onion and garlic: (34,37,40,41), bile: (42–44), wine: (45–47)). We explored the contribution of all four ingredients to both planktonic and biofilm activity of Bald’s eyesalve to build a picture of their relative contributions. Planktonic activity appeared almost entirely attributable to garlic. However, tests against *S. aureus* Newman biofilms, grown in a synthetic wound model, showed garlic exhibited no antibacterial activity in this more clinically-relevant setting. In fact, no preparation which omitted any one ingredient possessed full activity in the biofilm assay. This confirms our previously published finding that Bald’s eyesalve anti-biofilm activity is contingent on the presence of all four ingredients (20).

Our results against planktonic cultures of *S. aureus* and *P. aeruginosa* align with a report by Fuchs *et al.* 2018, that garlic alone, and specifically allicin, accounts for the majority of the planktonic activity of Bald’s remedy. Allicin is a defensive compound that is converted from alliin by the enzyme alliinase, upon damage to the plant tissue. Allicin can kill a wide range of both Gram-negative and Gram-positive bacteria *in vitro* (32) due to the thiosulfinate group (– S(O)–S–) reacting with many cellular proteins in the pathogen (48). We therefore explored the role of allicin in Bald’s eyesalve in detail.

Crucially, we found that 12 batches of Bald’s eyesalve were able to elicit much greater anti-biofilm activity than purified allicin at comparable concentrations. Pure allicin at a concentration of 500 μg·ml^-1^ was able to reduce viable cell numbers by 2-3 logs, whereas Bald’s eyesalve batches containing 275 - 486 μg·ml^-1^ allicin were able to cause a 6-7 log drop and eradicate the biofilms. Even in planktonic tests, we found that the concentration of allicin in the MIC of these batches of Bald’s eyesalve was lower than the MIC of purified allicin. Together, these results clearly illustrate that in addition to allicin, other ingredients in Bald’s eyesalve contribute to its activity. This highlights the importance of the combination of ingredients.

The differences between our findings and those of Fuchs *et al.* are partly explained by the contrasts between bacterial tolerance to killing (i) in MHB *versus* synthetic wound fluid; and (ii) in planktonic culture *versus* established biofilms. This highlights the differences in antibiotic susceptibility often seen in host-mimicking media versus standard MHB (18,19,49). Further, the allicin concentrations found in our batches of Bald’s eyesalve were lower than that found in Fuchs *et al.* The highest allicin concentration we measured was 486 µg·ml^-1^, compared with Fuchs *et al.*’s 836 µg·ml^-1^.

We think it unlikely that this is due to the differences in quantification methods used as both are appropriate. We used HPLC, a method that has been widely used previously (33,50–53) and is accepted to produce consistent and accurate results. Previous papers using HPLC have detected allicin concentrations ranging from 1 µg·ml^-1^ to 2 mg·ml^-1^ (33,52), a range that the results in this paper lie within. Fuchs *et al.* 2018 used quantitative NMR (qNMR), which too has advantages and is being increasingly used in quantification of concentrations (54). It is more likely that our Bald’s eyesalve preparations really do have different allicin concentrations. Alliin is released from storage vesicles and converted to allicin upon damage to tissue: Fuchs *et al.*, 2018 used a food processor, in comparison to our pestle and mortar, which may have resulted in greater damage to the tissue and therefore greater release of alliin and more allicin created. Further, different garlic varieties are thought to have different activities and compositions (55–59) and our research labs are based in different continents, which presumably results in different garlic varieties and growth conditions. The higher allicin concentration may explain the slightly increased planktonic activity seen in Fuchs *et al*. This may also be due to their use of agitation in MIC assays, as agitation is known to lower the MIC (60).

The step in the early Medieval remedy which specifies that the onion and garlic be ground together would have been conducted manually – most likely with a pestle and mortar. We have found that this process generates sub-bactericidal concentrations of allicin, which is complemented or synergised by the presence of other ingredients to form a biofilm-killing preparation. This demonstrates how contingent explorations of natural product antimicrobials are on material preparation and testing conditions. This result is also interesting because allicin can be toxic in high concentrations (61). Reducing the amount of allicin present but maintaining full activity against bacterial biofilms might have been key to producing a safe topical treatment. Further work is needed to elucidate the exact combination of natural products responsible for the anti-biofilm activity.

Contrary to many papers in the literature (45,62,63), we found that wine possessed very limited planktonic antimicrobial activity, and could not kill *S. aureus* in biofilms. Despite this, its absence from the full remedy causes a large drop in activity against *S. aureus* biofilms. Taken together, these observations suggest that the role of wine in the full recipe may be more to do with its physical properties, perhaps its ethanol content and/or low pH. Ethanol is a well-known extraction solvent and may allow better extraction of compounds from the plant matter (64), or allow better diffusion through biofilms, while the lower pH may activate pH-dependent compounds. The answer may lie in the combination of these properties, as the activity of wine reported in the literature cannot be attributed exclusively to one aspect of it: the physicochemical environment plays a large role in the activity of wine (65). Although pH and ethanol concentrations corresponding to those found in various wines have minimal effects on pathogens, without them the compounds within the wine have reduced activity (65).

Similarly, individual preparations of onion and bile, prepared as they would be for Bald’s eyesalve, possessed no planktonic antibacterial activity. This is contrary to the literature that has shown these ingredients to have activity against various strains (42,66). The most likely cause for this disparity is the difference in the preparation or the dilution effect when combined with other ingredients. However, the removal of onion or bile significantly reduced the anti-biofilm activity against *S. aureus*, indicating the importance of their presence within the whole remedy. They may be providing additional compounds necessary for the full killing effect in biofilms or aiding penetration/activity of antibacterial compounds from other ingredients.

Here, we have shown for the first time that Bald’s eyesalve has anti-biofilm activity against several clinically relevant bacteria, and we confirm our earlier work which concluded that this activity is dependent on combining all the ingredients. Interestingly, statistical analysis of the surviving portion of Bald’s Leechbook (London, British Library, Royal MS 12 D XVII) showed that garlic is combined with a second *Allium* species significantly more often than would be expected given the frequency of use of either garlic or other *Allium* species across the book if ingredients were combined randomly (67). Perhaps the patterns of ingredient combinations used by pre-modern physicians do, in at least some cases, reflect a requirement for combinatorial activity of several natural products to produce an efficacious antimicrobial preparation.

Research into natural products is often focussed on isolating single compounds, however, here we provide evidence that in doing so potent anti-biofilm mixtures can be overlooked. Viewing natural products in this way has the potential to open a vast new source of antimicrobials that can overcome the inherent difficulties of treating biofilm infections. In a companion manuscript (22), we describe the low potential of Bald’s eyesalve for producing irritation or impeding wound healing. Future work will determine whether our results translate into a candidate natural product cocktail for incorporation into wound ointments or dressings. At present, we conclude by re-stating the exciting potential for pre-modern European medical texts to contain antibacterial preparations of clinical interest.

## Acknowledgements

We thank Mathew Diggle, Kim Hardie, Ruth Massey and Tim Sloan for bacterial strains; Meera Unnikrishnan for helpful comments on an early draft of the manuscript; Callum Parsons, Colman Ó Cathail, Jason Millington and Thorulf Vargsen for pilot work; Callum Parsons, Jenny Littler, Shanjini Subhaskaran, Jason Millington, Thorulf Vargsen, Esther Sweeney and Colman O’Cathail for preparation of the eyesalve batches summarised in Figure 1; and our colleagues who helped initiate work on this project – Steve Diggle, Aled Roberts, Kendra Rumbaugh and Rebecca Gabrilska. We also thank Cerith Harries, Caroline Stewart and the University of Warwick Media Preparation team for preparing the media used for this work. This work was supported by a Diabetes UK Project Grant to FH (ref. 17/0005690) and Jessica Furner-Pardoe is funded by the MRC Doctoral Training Partnership [grant number MR/N014294/1].

## Supplementary Figures

**Figure S1.**
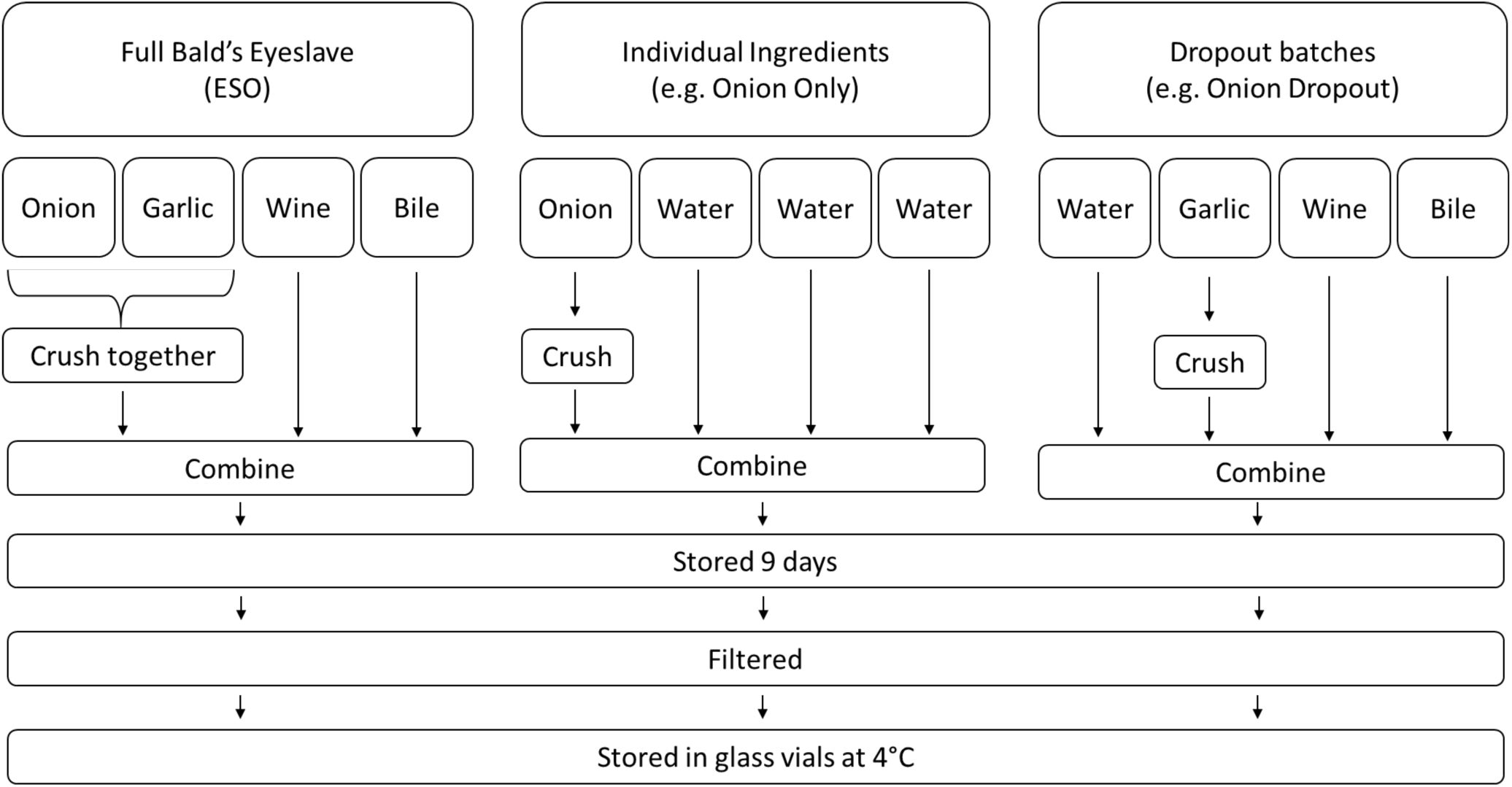
Schematic of the process to generate Bald’s eyesalve and various batch variations.

**Figure S2.**
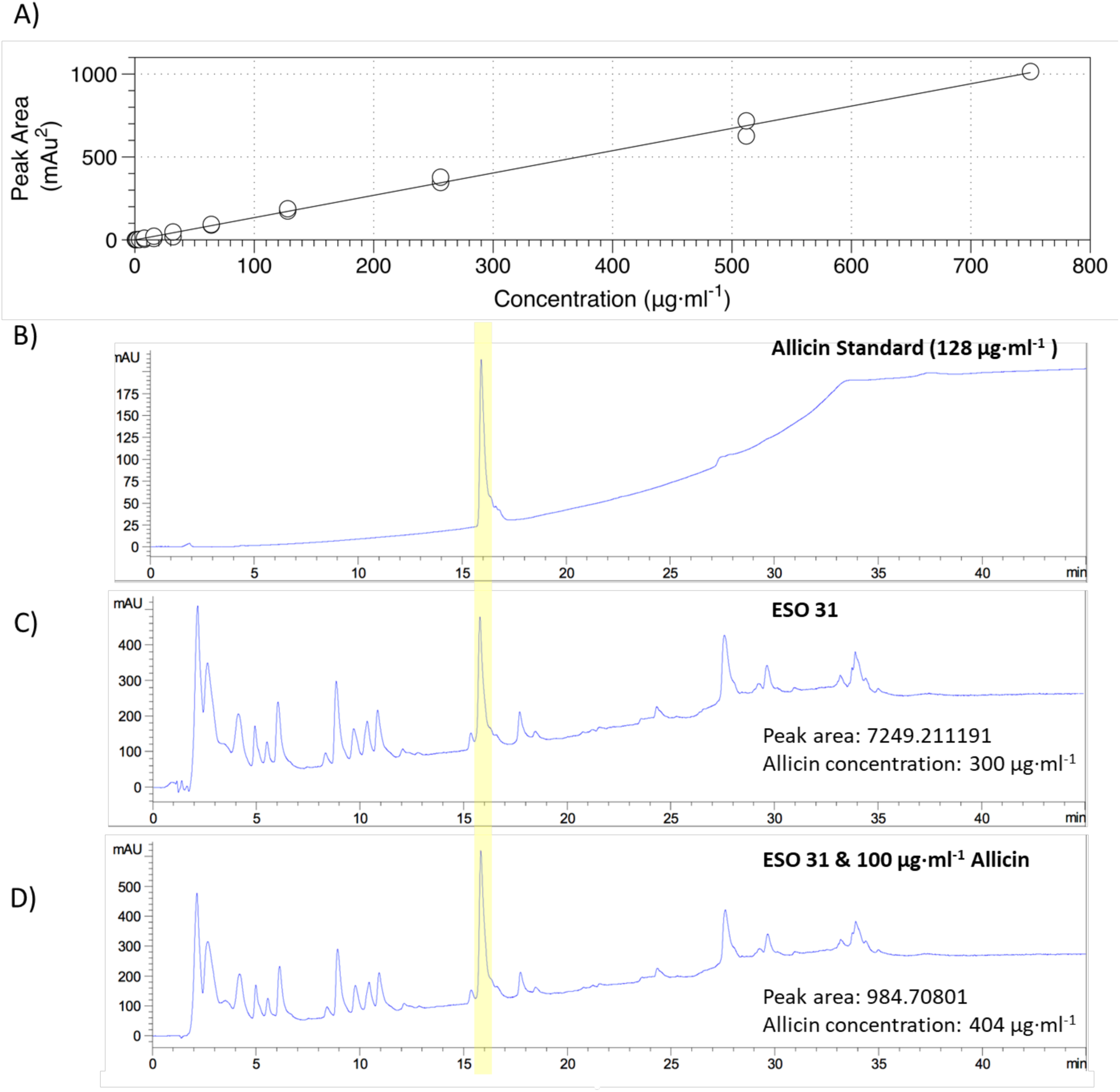
Confirmation of allicin in Bald’s eyesalve. All chromatograms were run against a 0-95% methanol gradient, with a flow rate of 1 ml·min^-1^ on HPLC Aligent 1200 series, at 210 nm UV wavelength. A) Allicin calibration curve. External allicin standards, serially diluted in water were run and peak area was measured. A linear relationship was found between HPLC peak area and allicin concentrations between 8 - 750 μg·ml^-1^ with a correlation coefficient R^2^ value > 0.99. The mid-range of the curve (8-512 μg·ml^-1^) has 2 repeats. B) Exemplary allicin external standard (128 μg·ml^-1^) chromatogram, suspected allicin peak has a retention time of approx. 15 minutes. C-D) Bald’s eyesalve fresh batch, ESO 31 without (C) and with (D) an additional 100 μg·ml^-1^ of allicin standard; the differences in peak area are indicated. The highlighted region is the allicin peak.

**Figure S3.**
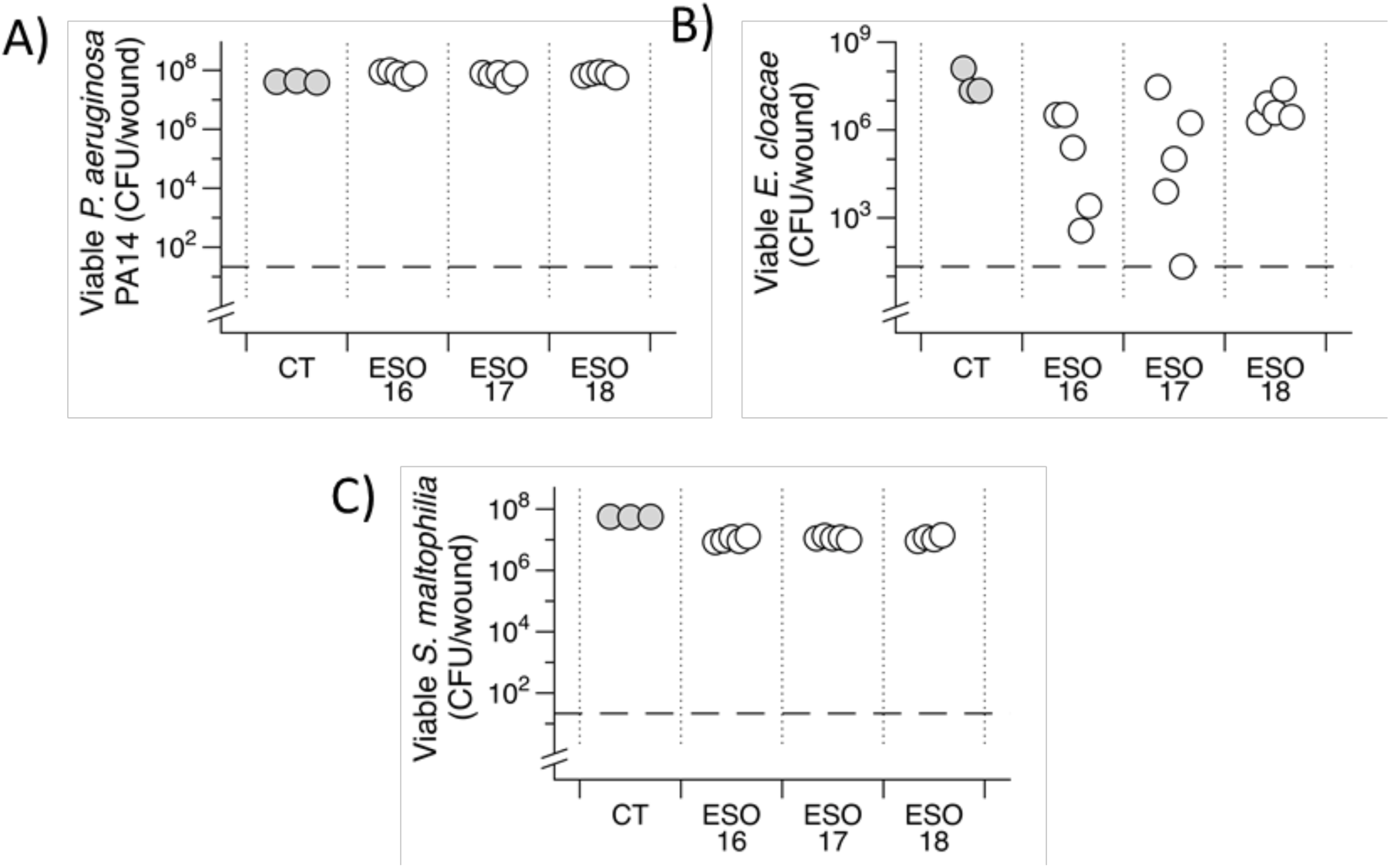
Bald’s eyesalve (ESO) anti-biofilm activity. Mature biofilms of various isolates were grown in a model of a soft tissue wound, then treated with either 0.5 vols sterile water (control, CT) or 0.5 vols of ESO for 24h before recovering bacteria for CFU counts (n = 3-5 replicates per treatment). The dashed line represents the limit of detection by plating. Data could not be transformed to fit assumptions of parametric tests, therefore, Kruskall-Wallis tests were used to determined that the CFU recovered from ESO-treated wounds was not significantly different from control wounds for *P. aeruginosa* PA14 (*X*^2^ = 6.88, df = 3, *p* value = 0.076), *S. maltophilia* (*X*^2^ = 7.63, df = 3, *p* value = 0.54) or *E. cloacae* (*X*^2^ = 7.74, df = 3, *p* value = 0.051). Raw data and R scripts are supplied in the Data Supplement.

## Notes

### Competing Interest Statement

The authors have declared no competing interest.

### Summary of Updates

DOI added for Anonye et al. Reference list copyedited for clarity.

